# Activation of EGFR signaling by Tc-Vein and Tc-Spitz regulates the metamorphic transition in the red flour beetle *Tribolium castaneum*

**DOI:** 10.1101/2021.04.20.440594

**Authors:** Sílvia Chafino, David Martín, Xavier Franch-Marro

## Abstract

Animal development relies on a sequence of specific stages that allow the formation of adult structures with a determined size. In general, juvenile stages are dedicated mainly to growth, whereas last stages are devoted predominantly to the maturation of adult structures. In holometabolous insects, metamorphosis marks the end of the growth period as the animals stops feeding and initiate the final differentiation of the tissues. This transition is controlled by the steroid hormone ecdysone produced in the prothoracic gland. In Drosophila different signals have been shown to regulate the production of ecdysone, such as PTTH/Torso, TGFß and Egfr signaling. However, to which extent the role of these signals is conserved remains unknown. Here, we study the role of Egfr signaling in post-embryonic development of the basal holometabolous beetle *Tribolium castaneum*. We show that *Tc-Egfr and Tc-pointed* are required to induced a proper larval-pupal transition through the control of the expression of ecdysone biosynthetic genes. Furthermore, we identified an additional Tc-Egfr ligand in the *Tribolium* genome, the neuregulin-like protein Tc-Vein (Tc-Vn), which contributes to induce larval-pupal transition together with Tc-Spitz (Tc-Spi). Interestingly, we found that in addition to the redundant role in the control of pupa formation, each ligand possesses different functions in organ morphogenesis. Whereas Tc-Spi acts as the main ligand in urogomphi and gin traps, Tc-Vn is required in wings and elytra. Altogether, our findings show that in *Tribolium*, post-embryonic Tc-Egfr signaling activation depends on the presence of two ligands and that its role in metamorphic transition is conserved in holometabolous insects.

## INTRODUCTION

Animal development consists of a number of stage-specific transitions that allows the growth of the organism and the proper morphogenesis of adult structures. Whereas growth is restricted to juvenile stages, final adult differentiation and sexual maturity takes place without tissue proliferation. Thus, the transition from juvenile to adult stage determines the final size of the animal. Examples of this particular transition are puberty in humans and metamorphosis in insects. Despite its importance, however, the precise mechanisms underlying the regulation of this developmental transition are far from being clearly understood.

Holometabolous insects are a paradigm for the study of the precise control of the transition between the immature larva and the adult, which happens through a transitional metamorphic stage, the pupa. In these insects, a pulse of the steroid hormone 20-hydroxyecdysone (20E) at the end of the larval period triggers the onset of metamorphosis [1,2]. Biosynthesis of ecdysone, the precursor of 20E, takes place in a specialized organ called the prothoracic gland (PG) and is controlled by the restricted expression of several biosynthetic enzymes encoding genes collectively referred to as the *Halloween* genes. These include the Rieske-domain protein *neverland* (*nvd*) [3,4], the short-chain dehydrogenase/reductase *shroud* (*sro*) [5] and the P450 enzymes *spook* (*spo*), *spookier* (*spok*), *phantom* (*phm*), *disembodied* (*dib*) and *shadow* (*sad*) [6–11]. Work carried out in the fruit fly *Drosophila melanogaster* revealed that the proper function of the PG at the metamorphic transition, including the timely expression of the *Halloween* genes, involves the integrated activity of different signaling pathways such as TGFß/activin [12], Insulin receptor (InR) [13–16], and PTTH/Torso pathways [17–19]. Whereas insulin and PTTH/Torso pathways promote the up-regulation of the *Halloween* genes at the appropriate developmental time to trigger metamorphosis [2], TGFß/activin signaling regulates insulin and Ptth/torso pathways in the PG by controlling the expression of their respective receptors [12]. In addition to these pathways, we have recently shown that the activation of the Epidermal Growth Factor Receptor (Egfr) signaling pathway in PG cells is also critical for ecdysone production at the metamorphic transition [20]. Thus, Egfr pathway activation controls *Halloween* gene expression and ecdysone vesicle secretion during the last larval stage. Importantly, it has been shown that the control of the metamorphic transition by TGFß/activin, Insulin receptor (InR), and PTTH/Torso pathways is conserved in less derived holometabolous insects [21–28]. However, the grade of conservation of the Egfr signaling pathway in this process has not been well established.

In *Drosophila*, Egfr is activated by four ligands with a predicted Egf-like motif: the TGF-like proteins Gurken (Grk), Spitz (Spi) and Keren (krn), and the neuregulin-like protein Vein (Vn). The specific expression and activity of these ligands in different tissues appear to be responsible for different levels of EGF signaling activation in particular developmental contexts [29,30]. For example, Krn acts redundantly with Spi in the embryo, during eye development and in adult gut homeostasis [31–33], whereas Grk is specific of the germline, where it acts to establish egg polarity [34,35]. In contrast, Vn acts as the main ligand in the wing, in muscle attachment sites, during the air sac primordium development and in the patterning of the distal leg region [36–40]. Once Egfr is activated by any of its ligands, the signal is relayed through the sequential activation of the MAPK/ERK kinase pathway to the nucleus, where is finally mediated by the transcription factor pointed (pnt) [41,42].

In the present work, we aim to study the role of Egfr signaling pathway in the control of the metamorphic transition in the more basal holometabolous insect, the red flour beetle *Tribolium castaneum*, which diverged from *Drosophila* ∼ 250 million years ago. Despite the early divergence between both species, the role of this pathway in the control of several developmental processes has been revealed to be conserved. For example, it induces the encapsulation of the oocyte by the somatic follicle cell layer and establishes the polarity of the egg chambers and the D-V axis of the embryo during oogenesis [43]. It also controls the formation and patterning of the legs and the proper development of the abdomen and Malpighian tubules during embryogenesis [43–45], and regulates distal development of most appendages such as leg, antenna, maxilla and labium, and promotes axis elongation of the mandibles [46–48]. Despite this functional conservation, a remarkable difference between *Drosophila* and *Tribolium* is found in the number of Egfr ligands identified, for only a single TGF-EGF ligand, Tc-Spi, has been found in the beetle [44].

In here, we confirm that Tc-Egfr signaling is required for a normal ecdysone-dependent transition from larval to pupal stages in *Tribolium*. We found that inactivation of Tc-Egfr signaling in the last larval stage results in the arrest of larval development at the larval-pupal transition, with arrested larvae presenting reduced levels of the Halloween gene *Tc-phm*, as well as of the ecdysone-dependent transcription factors *Tc-Hr3, Tc-E75* and *Tc-Broad Complex* (*Tc-Br-C*). Importantly, we provided evidence of the existence of an additional Tc-Egfr ligand in *Tribolium*, the neuregulin-like protein Tc-Vein (Tc-Vn). We show that Tc-Vn and Tc-spi act redundantly in the control of larval-pupal transition, while function separately during pupal morphogenesis. Taken together, our results strongly suggest that Egfr signaling pathway plays a conserved central role in the control of the ecdysone-dependent larval-pupal transition and during the morphogenesis of the pupa in holometabolous insects.

## 3. RESULTS

### Egfr signaling is required for larval-pupal transition in *Tribolium*

As a first step towards the characterization of Egfr signaling in post-embryonic *Tribolium*, we measured mRNA levels of *Tc-Egfr* and *Tc-pnt* by RT-qPCR in staged penultimate (L6) and last (L7) instar larvae. Expression of *Tc-Egfr* and *Tc-pnt* was low at the onset of both instars, and then steadily increased to reach the maximal level at the final part of each instar, which suggests a role of Egfr signaling during stage transitions (Fig 1A). To examine this possibility, we analyzed the functions of both factors depleting *Tc-Egfr* and *Tc-pnt* by injecting dsRNAs for each transcript in L6 instar larvae (*Tc-Egfr*^*RNAi*^ *and Tc-pnt*^*RNAi*^ animals). Specimens injected with *dsMock* were used as negative controls (*Control* animals). All *Tc-Egfr*^*RNAi*^ *and Tc-pnt*^*RNAi*^ L6 larvae molted to normal L7 larvae but then failed to pupate at the ensuing molt, as did *Control* larvae. Instead, they arrested development during the larval-pupal transition (Fig 1B). The nature of the affected transition in the arrested animals, from larva to pupa, was analyzed by measuring the expression levels of the metamorphosis-triggering factor *Tc-E93* [49]. As Fig. 1C shows, arrested *Tc-Egfr*^*RNAi*^ and *Tc-pnt*^*RNAi*^ animals showed normal expression of *Tc-E93* thus indicating that lack of Egfr signaling did not affect the nature of the larval-pupal transition (Fig 1C).

**Figure 1.**
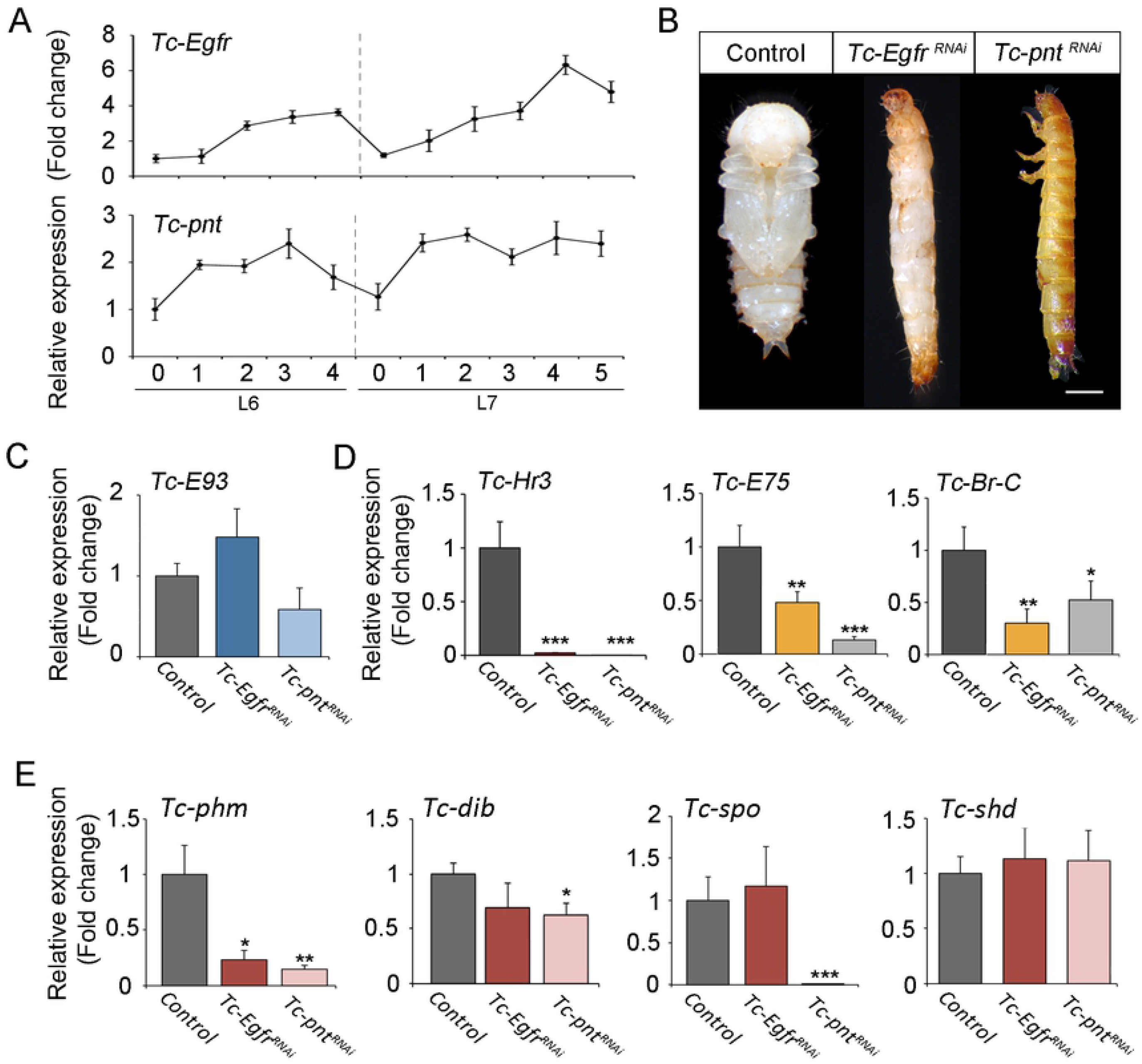
Egfr signalling controls larval-pupal transition in *Tribolium*. (A) *Tc-Egfr* and *Tc-pnt* mRNA levels measured by qRT-PCR in penultimate (L6) and ultimate (L7) instar larvae. Transcript abundance values are normalized against the *Tc-Rpl32* transcript. Fold changes are relative to the expression of *Tc-Egfr* and *Tc-pnt* in day 0 of L6 larvae, arbitrarily set to 1. Error bars indicate the SEM (n=5). (B) L6 larvae were injected with *dsMock* (*Control*) or with *dsTc-Egfr* (*Tc-Egfr*^*RNAi*^), and *dsTc-pnt* (*Tc-pnt*^*RNAi*^) and left until the ensuing molts. Ventral views of a *Control* pupa and *Tc-Egfr*^*RNAi*^ and *Tc-pnt*^*RNAi*^ larvae arrested at the larval-pupal transition. Scale bar represents 0.5mm. (C-E) Transcript levels of *Tc-E93* (C), *Tc-Hr3, Tc-E75* and *TcBr-C* (D) and biosynthetic ecdysone genes *Tc-phm, Tc-dib, Tc-spok and Tc-shd* measured by qRT-PCR in 5-day-old L7 *Control, Tc-Egfr*^*RNAi*^, and *Tc-pnt*^*RNAi*^ larvae. Transcript abundance values were normalized against the *Tc-Rpl32* transcript. Average values of three independent datasets are shown with standard errors. Asterisks indicate differences statistically significant at * p<0.05 and ** p<0.005 (t-test).

Since *Tc-Egfr* and *Tc-pnt* were also expressed in L6, we studied whether Egfr signaling was also required in earlier larval-larval transitions. To this aim, we injected dsRNA of *Tc-Egfr* in antepenultimate L5 larvae. Under these conditions, all L5-*Tc-Egfr*^*RNAi*^ larvae developed normally and underwent two successive molts until reaching L7, when larvae arrested development before pupation confirming that Egfr signaling is only required for the last larval transition.

Since the larval-pupal transition is ecdysone-dependent, the phenotype presented by *Tc-Egfr*^*RNAi*^ *and Tc-pnt*^*RNAi*^ larvae is consistent with an ecdysone-signaling deficiency. To assess this hypothesis, we measured mRNA expression levels of the direct ecdysone-dependent genes *Tc-Hr3, Tc-E75* and *Tc-Br-C* in late L7 larvae, which are commonly used as proxies for ecdysone levels [17,50,51]. As Figure 1D shows, *Tc-Egfr*^*RNAi*^ *and Tc-pnt*^*RNAi*^ larvae presented significantly reduced expression levels of these genes compared to *Controls*. Consistently, the expression of the *Halloween* gene *phm* was also decreased in *Tc-Egfr*^*RNAi*^ *and Tc-pnt*^*RNAi*^ larvae (Fig 1E). However, the expression of three other *Halloween* genes, *Tc-dib, Tc-spook* and *Tc-shd*, was not affected by the absence of *Tc-Egfr* (Fig 1E), indicating that the downregulation of *Tc-phm* is specific and not due to a general transcriptional effect. Interestingly, in addition to the transcriptional regulation of *Tc-phm*, the gene expression levels of *Tc-dib* and *Tc-spok* were also downregulated in *Tc-pnt*^*RNAi*^ larvae compared to *Controls* (Fig 1E), indicating a stronger inactivation of Tc-Egfr signalling upon depletion of this transcription factor. Taken together, these results indicate that Tc-Egfr signaling is specifically required for a proper ecdysone-dependent larval-pupal transition *in Tribolium*.

### Identification of the Tc-Egfr ligand Tc-Vein in *Tribolium*

The next question was to determine which EGF ligand was responsible for the activation of the Tc-Egfr pathway during the larval-pupal transition. As stated before, the TGF∼-like protein Tc-Spi is the only Tc-Egfr ligand identified in *Tribolium* [44]. Expression analysis of *Tc-spi* during the last two larval instars revealed that it is expressed in a similar pattern than that of *Tc-Egfr* (Fig 2A), which suggests that Tc-Spi might act as the Tc-Egfr ligand during the transition. We studied this possibility by injecting *dsTc-spi* into L6 larvae (*Tc-spi*^*RNAi*^ animals). Unexpectedly, and in contrast to *Tc-Egfr*^*RNAi*^ *and Tc-pnt*^*RNAi*^ larvae, all L7-*Tc-spi*^*RNAi*^ larvae molted to L7 and then to pupa on a normal schedule (Fig 2B), suggesting the occurrence of different Tc-Egfr ligands during the post-embryonic development of *Tribolium*.

**Figure 2.**
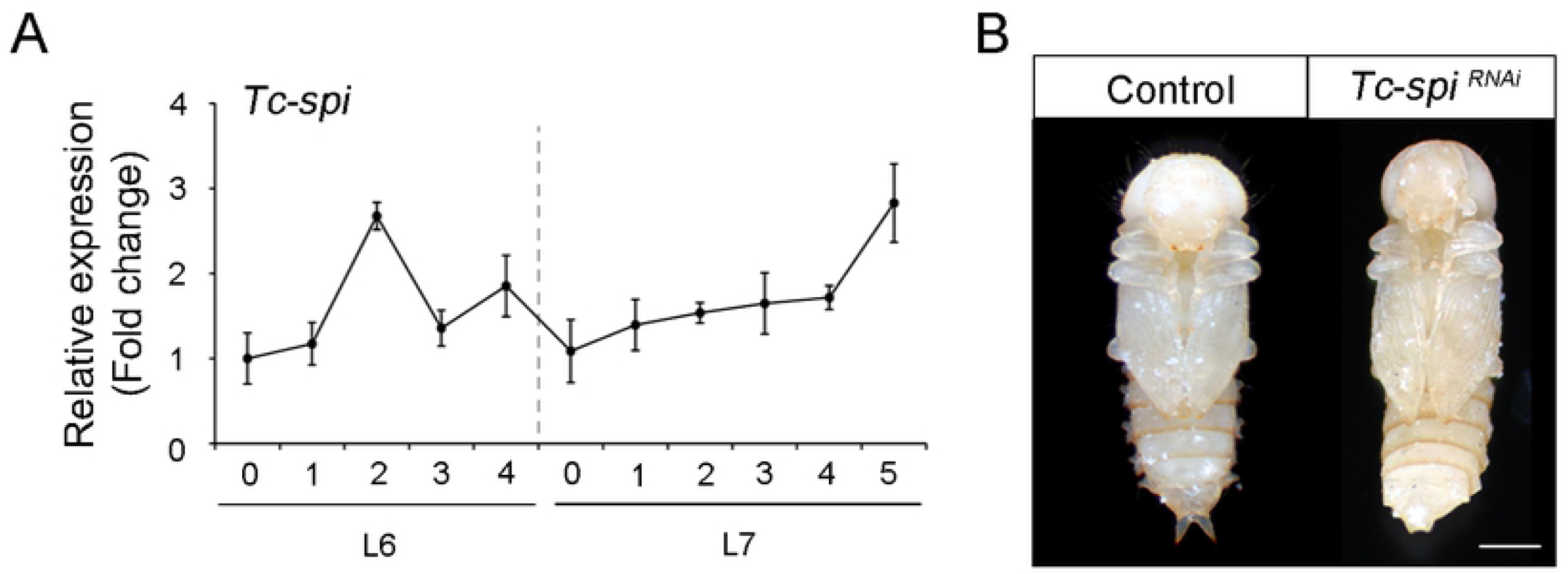
Tc-Spi is not the solely ligand of Tc-Egfr signaling during larval-pupal transition in *Tribolium*. (A) *Tc-spi* mRNA levels measured by qRT-PCR in penultimate (L6) and ultimate (L7) instar larvae. Transcript abundance values are normalized against the *Tc-Rpl32* transcript. Fold changes are relative to the expression of *Tc-Egfr* and *Tc-pnt* in day 0 of L6 larvae, arbitrarily set to 1. Error bars indicate the SEM (n=5). (B) L6 larvae were injected with *dsMock* (*Control*) or with *dsTc-spi* (*Tc-spi*^*RNAi*^) and left until the ensuing molts. Ventral views of *Control* and *Tc-spi*^*RNAi*^ pupae. Scale bar represents 0.5mm.

In *Drosophila* four different Egf ligands have been identified: Dm-Grk, Dm-Ker and Dm-Spi are TGF-like proteins, whereas Dm-Vn belongs to the neuregulin-like family [29]. Tc-Spi is highly similar to the three *Drosophila* genes Dm-Grk, Dm-Spi and Dm-Ker [31,52]. In contrast, no neuregulin-like protein has been identified in *Tribolium*. To investigate the presence of a neuregulin-like Egf ligand, we performed a detailed Tblastn search in the beetle genome database with the Dm-Vn sequence of *Drosophila*. This search revealed the presence of a Tc-Vn orthologue in *Tribolium*. The predicted Tc-Vn protein has 380 amino acids and presents conserved EGF-like and neuregulin Ig-like domains as well as a PEST region, all of which are characteristic of Vn proteins [53] (Fig 3A and S1 Fig). Additionally, Tc-Vn contains a hydrophobic region at the amino terminus that is typical of a signal sequence, a feature that is consistent with Tc-Vn being a secreted protein. The neuregulin Ig-like domain consists of 100 amino acids and presents the two invariant cysteine residues typical of the domain (Fig 3B and S1 Fig). The EGF-like domain is 47 amino acids long and presents the six invariant cysteines and highly conserved glycine and arginine residues characteristic of the motif (Fig. 3C and Supplementary Fig. 1; [54,55]). Finally, Tc-Vn contains a 24 amino acid long PEST region, characteristic of proteins with short half-lives, that localizes between the EGF-like and neuregulin Ig-like domains (Fig. 3A and Supplementary Fig. 1). Phylogenetic analysis of Vn protein sequences showed that Tc-Vn grouped with others coleopteran sequences, confirming that Tc-Vn belongs to neuregulin Ig-like family (Fig. 4).

**Figure 3.**
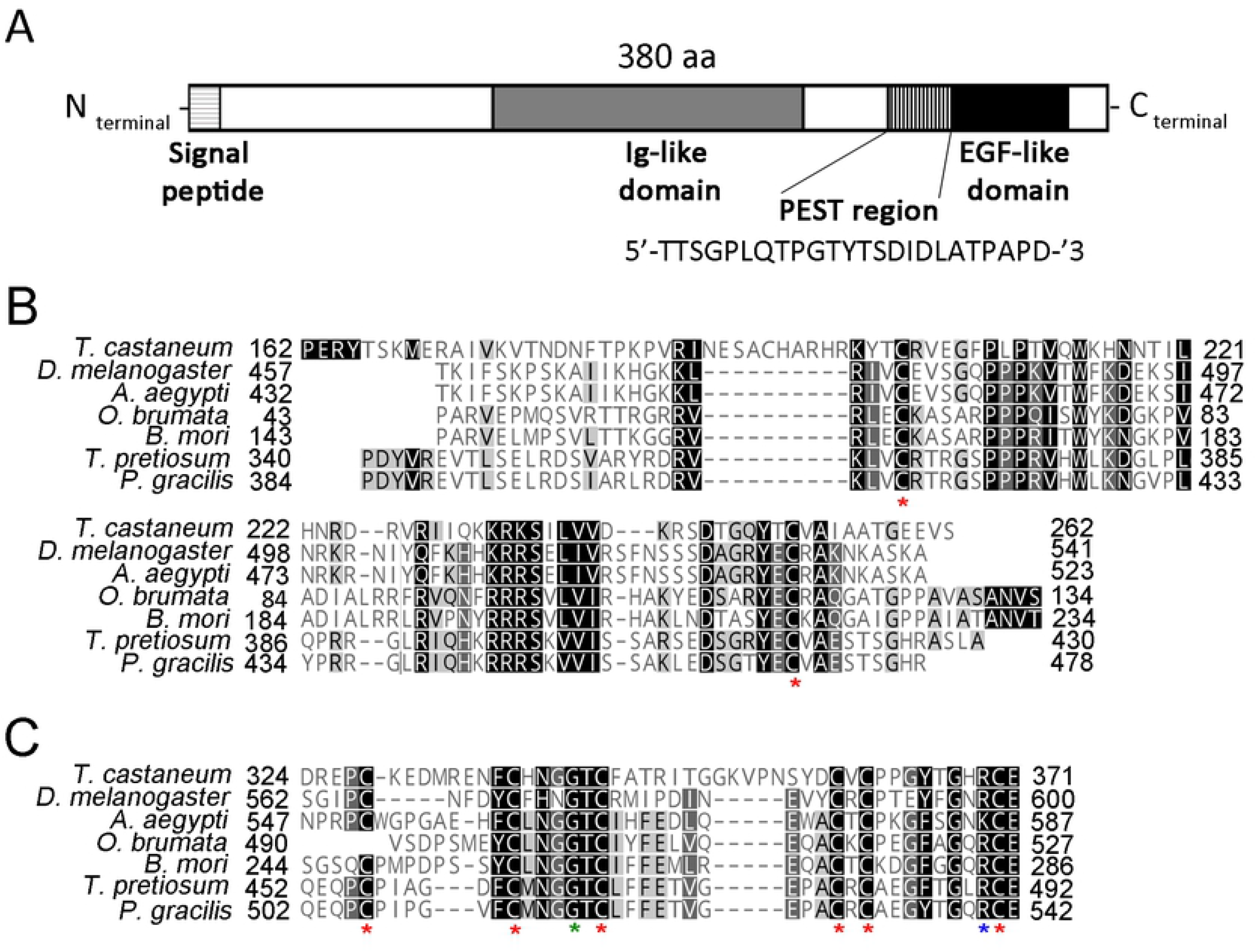
The structure of *Tribolium* Tc-Vn protein. (A) Predicted amino acid sequence of Tc-Vn. The signal peptide is shown in horizontal striped box. Two highly conserved protein domains Ig-like (grey box) and EGFR-like (black box) are indicated. The PEST region is represented by vertical striped box. (B) Protein alignment of Ig-like domain and C) Egfr-like domain. Asterisks in red indicate the invariant highly conserved cysteines, in green the conserved Glycine and in blue the conserved Arginine from IG-like and Egfr-like domains.

**Figure 4.**
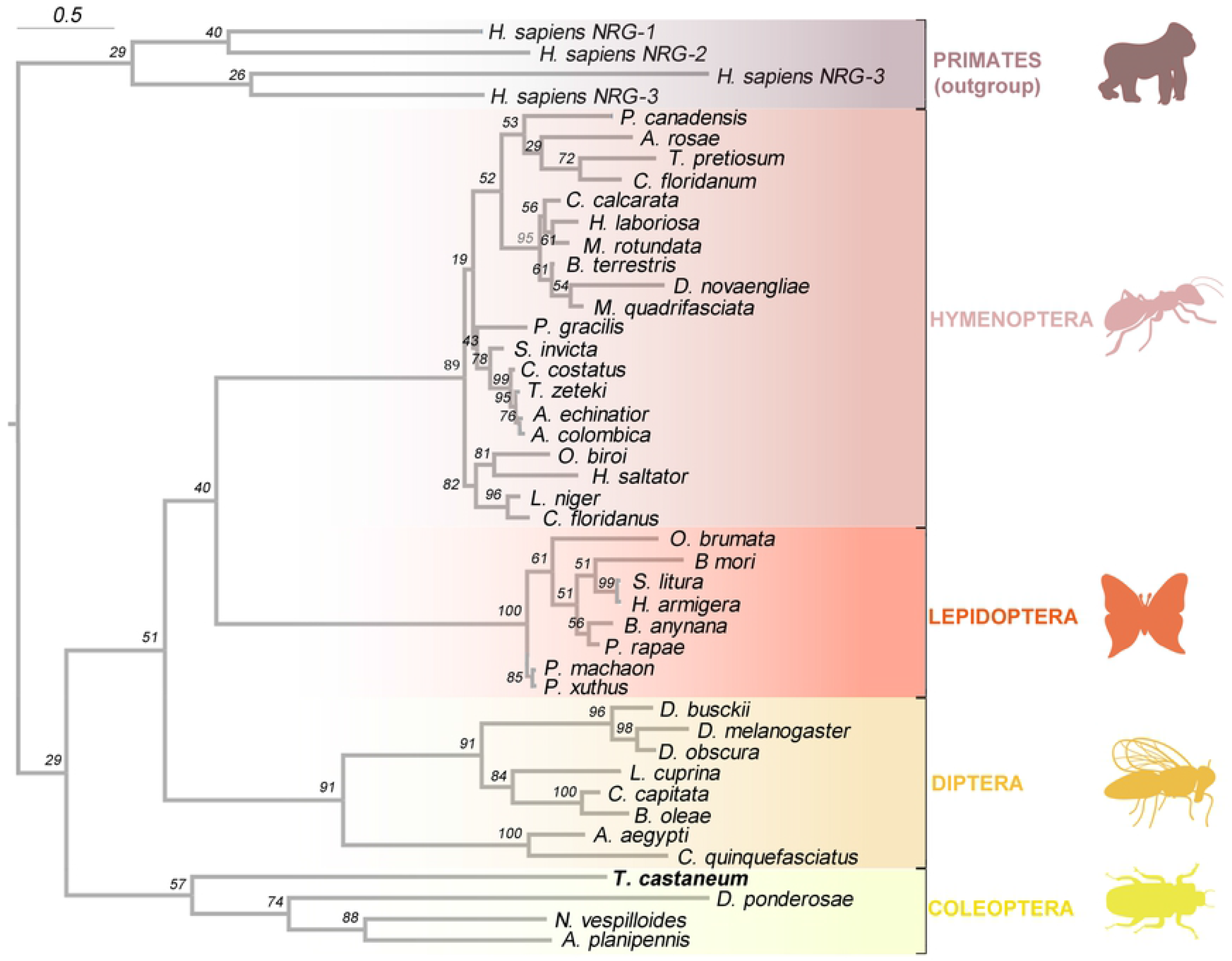
Phylogenetic analysis of Vn proteins. Phylogenetic tree based in Vn protein sequences from 44 different insect taxa including the Tc-Vn sequence of *Tribolium* described in this study. Four primate Hs-Vn sequences are used as outgroup. *Tribolium* is shown in red. The different insect orders are indicated in right part of the phylogenetic tree.

### Tc-Vn and Tc-Spi have redundant functions in larval-pupal transition

Since Tc-Vn is a newly identified protein, we wanted to determine the role of this ligand in *Tribolium*. Tc-Egfr signaling has been already involved in the regulation of key processes in embryonic, metamorphic and adult stages [43–48]. Thus, parental depletion of *Tc-Spi, Tc-Egfr*, and *Tc-Pnt* severely reduced egg production, and also resulted in embryos with shorter appendages and problems in the development of thoracic and abdominal segments [43,44]. To study the function of Tc-vn in adult and embryonic stages, we injected *dsTc-vn* into adult females (*Tc-vn*^*RNAi*^ animals) and found that, in contrast to what is observed in *Tc-Egfr*-depleted animals, their ovaries developed properly, and eggs were laid normally (S2A-D Fig). Likewise, the resulting embryos developed as normal and eclosed on a regular schedule (data not shown). Consistent with this finding, whereas the relative expression of *Tc-spi* was strongly detected in ovaries and embryos the levels of Tc-vn were almost neglected, supporting the idea that *Tc-vn* is dispensable for Tc-Egfr activation in adult and embryonic stages (S2E Fig).

To characterize Tc-Vn during post embryonic stages we next analyzed its expression pattern during the last two larval instars. As Fig. 6A shows, *Tc-vn* is up-regulated at the first day of L6 and its expression is maintained until the beginning of L7 when it declined. Then, *Tc-vn* is up-regulated again to reach the highest levels in the prepupal stage similarly to *Tc-spi* expression. To ascertain the role of the ligand, we next injected *dsTc-vn* in L6 larvae. Similar to the *Tc-spi*^*RNAi*^ phenotype, L6-*Tc-vn*^*RNAi*^ animals molted to normal L7 larvae and then to pupae although with some morphological defects (Fig 5B and Fig 6A’-F’). These results demonstrate that Tc-Vn is able to activate the Egfr signaling pathway during the larval-pupal transition.

**Figure 5.**
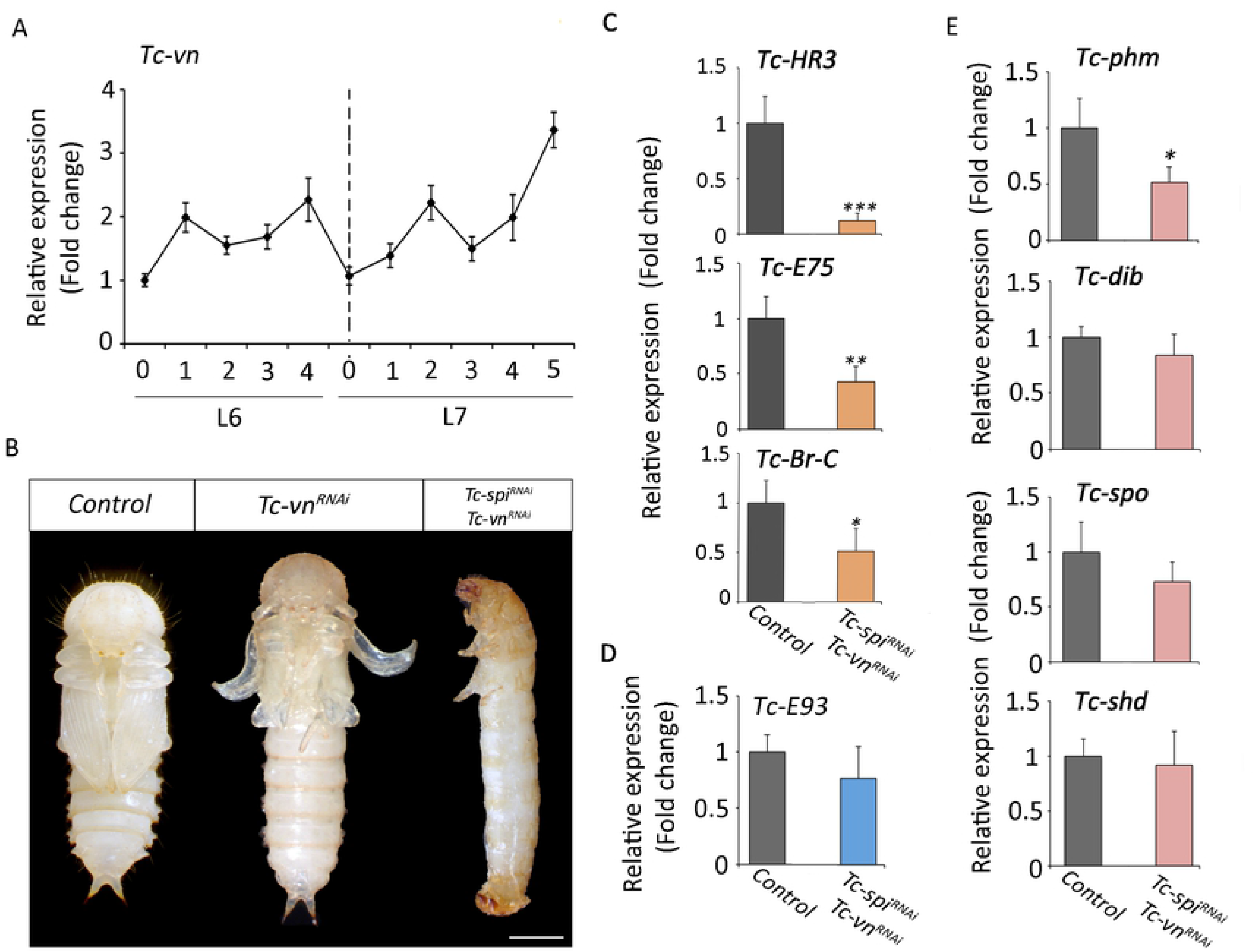
Tc-Vn and Tc-Spi activate redundantly the larval-pupal transition in *Tribolium*. (A) *Tc-vn* mRNA levels measured by qRT-PCR in penultimate (L6) and ultimate (L7) instar larvae. Transcript abundance values are normalized against the *Tc-Rpl32* transcript. Fold changes are relative to the expression of *Tc-vn* in day 0 of L6 larvae, arbitrarily set to 1. Error bars indicate the SEM (n=5). (B) L6 larvae were injected with *dsMock* (*Control*), *dsTc-spi* (*Tc-spi*^*RNAi*^) or both ds*Tc-spi* and *dsTc-vn* simultaneously (*Tc-spi*^*RNAi*^ + *Tc-vn*^*RNAi*^) and left until the ensuing molts. Ventral views of *Control* and *Tc-spi*^*RNAi*^ pupae and a *Tc-spi*^*RNAi*^ + *Tc-vn*^*RNAi*^ larva arrested at the larval-pupal transition. Scale bar represents 0.5 mm. (C) Transcript levels of *Tc-Hr3, Tc-E75* and *TcBr-C*, (D) *Tc-E93* and (E) biosynthetic ecdysone genes *Tc-phm, Tc-dib, Tc-spok and Tc-shd* measured by qRT-PCR in 5-day-old *Control*, and *Tc-spi*^*RNAi*^ + *Tcvn*^*RNAi*^ L7 larvae. Transcript abundance values were normalized against the *Tc-Rpl32* transcript. Average values of three independent datasets are shown with standard errors. Asterisks indicate differences statistically significant at * p<0.05 and ** p<0.005 (t-test) *** p<0.0005 (t-test).

**Figure 6.**
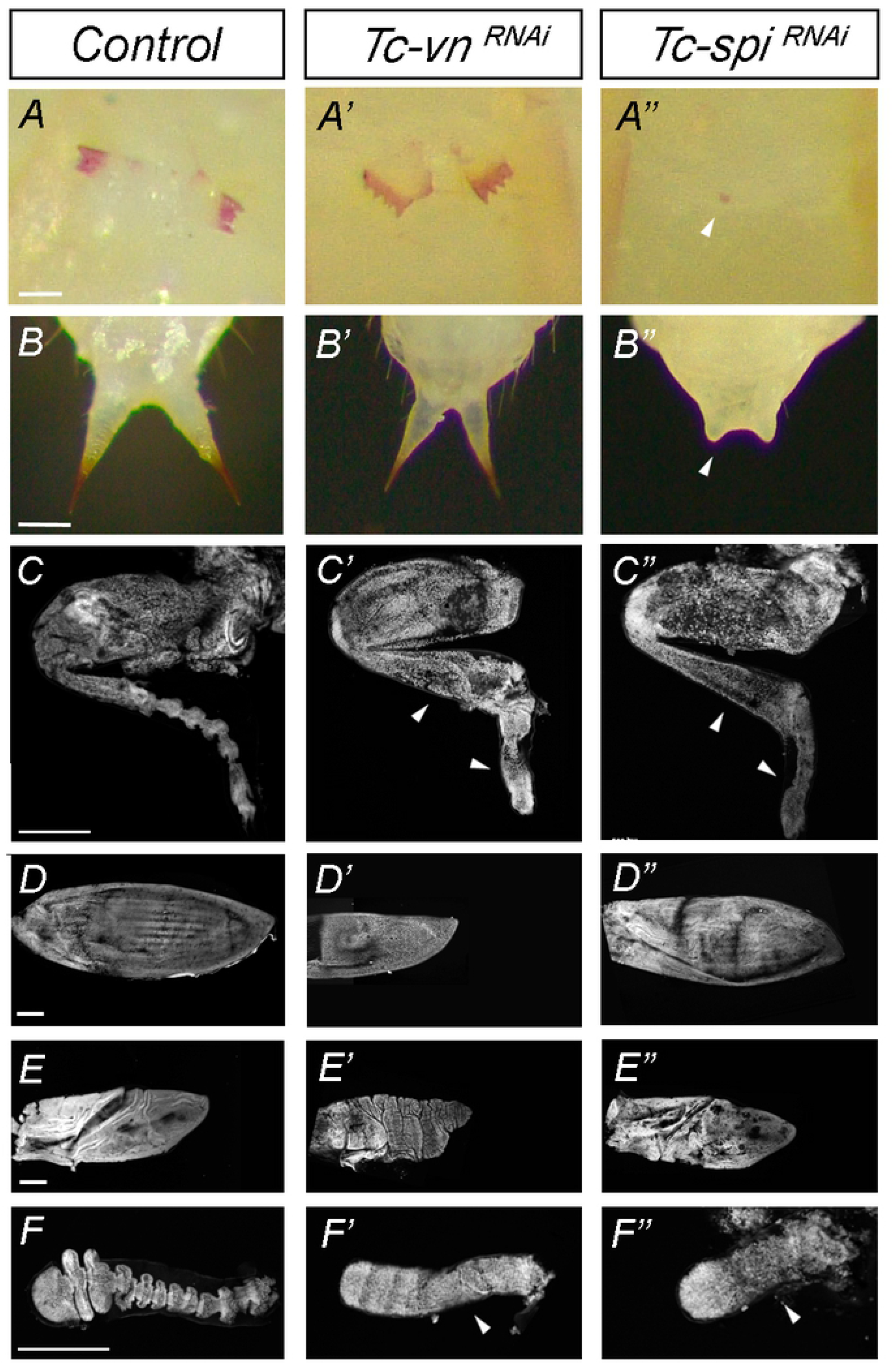
Differential role of Tc-Vn and Tc-Spi in the morphogenesis of pupal structures. Comparison of the external morphology of pupal appendages between (A-F) *Control*, (A’-F’) *Tc-vn*^*RNAi*^ and (A’’-F’’) *Tc-spi*^*RNAi*^ pupae. The pupal structures are (A-A’’) Gin traps, (B-B’’) urogomphi, (C-C’’) hindlegs, (D-D’’) elytra, (E-E’’) wing and (F-F’’) antenna. The scale bars represent 0.1 mm in A-A’’, 0.3 mm in B-B’’ and 500 ∼m in C-C’’, D-D’’, E-E’’ and F-F’’.

The fact that the depletion of neither *Tc-vn* nor *Tc-spi* prevents larval-pupal transition raises the possibility that both ligands might redundantly activate the Egfr signaling pathway at this stage of development. To address this question, we interfered *Tc-vn* and *Tc-spi* simultaneously in penultimate L6 larvae (*Tc-vn*^*RNAi*^+*Tc-spi*^*RNAi*^ animals). Interestingly, all *Tc-vn*^*RNAi*^+*Tc-spi*^*RNAi*^ animals molted to L7 but then arrested development at the end of the last larval stage, as *Tc-Egfr* or *Tc-pnt* larvae (Fig 5B). This observation was further confirmed by the analysis of *Tc-HR3, Tc-E75, Tc-Br-C* mRNA levels, which were strongly reduced in *Tc-vn*^*RNAi*^+*Tc-spi*^*RNAi*^ larvae compared to the *Control* (Fig 5C). Likewise, the expression of *Tc-E93* was not significantly affected indicating that depletion of both ligands does not influence to nature of molt as when the *Tc-Egf* receptor was depleted (Fig. 5D and Fig 1C). Furthermore, the expression of *Tc-phm* was also decreased in double knockdown animals whereas the expression levels of *Tc-dib, Tc-spok* and *Tc-shd* were not affected (Fig. 5E). Altogether, these results demonstrate that *Tc-vn* and *Tc-spi* can act redundantly in the control of the metamorphic transition in *Tribolium*.

### Tc-Spi and Tc-Vn control different aspects of *Tribolium* metamorphosis

As previously showed, *Tc-vn*^*RNAi*^ and *Tc-spi*^*RNAi*^ animals were able to pupate, although with morphological defects (compare Fig 2B and Fig 5B). Interestingly, some of those defects were ligand specific. For example, *Tc-spi*^*RNAi*^ pupae lacked gin traps and well developed urogomphi (Fig 6A’’-F’’), whereas *Tc-vn*^*RNAi*^ pupae showed normal gin traps and urogomphi but presented curved and smaller elytra and wings compared to *Tc-spi*^*RNAi*^ animals (Fig 6A’-E’). In contrast, both knockdown pupae showed similar morphological defects in the segmentation of the antenna and the distal part of the legs (Fig 6C-C’’ and 6F-F’’). These results suggest that Tc-Vn and Tc-Spi activate the Egfr signaling pathway in a tissue specific manner during the pupal transformation.

## DISCUSSION

Egfr signaling in insects is involved in the regulation of multiple processes during organism development such as cell survival, proliferation, and differentiation. In addition to these functions, it has been recently shown in *Drosophila* that regulates the metamorphic transition by controlling ecdysone biosynthesis in the PG [20]. Interestingly, our study suggests that this role is conserved in the more basal holometabolous insect *Tribolium*, as the activity of this pathway is specifically required for the larval-pupal transition in this beetle. Furthermore, our study revealed the existence of an additional EGF ligand in *Tribolium*, the neuroglianin Tc-Vn, which, together with the already described Tc-Spi ligand, activates Egfr in a redundant manner for the induction of the metamorphic transition. In contrast, we report different requirements of both ligands for the formation of pupal structures, such as wings, elytra or gin traps, suggesting a ligand-specific activation of Egfr signaling during metamorphosis.

### Egfr signaling regulates larval-pupal transition in *Tribolium*

In *Drosophila* Egfr signaling regulates larval-pupal transition through the control of PG size and the expression of ecdysone biosynthetic genes in this gland [20]. Our results in *Tribolium* suggest that such regulation might be general feature of holometabolous insects. Several evidences support this possibility: (*i*) depletion of both Tc-Egfr ligands, *Tc-vn* and *Tc-spi*, is required to induce the arrested development phenotype whereas individual depletion of each ligand results in pupa formation with specific morphological defects; (*ii*) depletion of the main transductor of the Egfr signaling *Tc-pnt* results in the same phenotype; (*iii*) the requirement of Egfr signaling is restricted to the last larval stage as larval-larval molting is unaffected upon inactivation of *Tc-Egfr* in early larval stages, and (*iv*) depletion of *Tc-Egfr* or its ligands results in reduced levels of the Halloween gene *Tc-phm* as well as of several 20E-dependent genes during the larva-pupa transition. However, we need to take into consideration that the use of systemic RNAi to inactivate Tc-Egfr signaling in *Tribolium* cannot discard an indirect effect on ecdysone production. In this regard, further studies in other holometabolous group of insects would help confirm this evolutionary conserved trait.

It is interesting to note that 20E controls not only the metamorphic switch but also all the previous larval-larval transitions. The fact that Egfr signaling is involved specifically in triggering pupa formation suggests that other factors might control the production of ecdysone in the previous transitions. In this sense, several transcription factors such as Ventral veins lacking (Vvl), Knirps (Kni) and Molting defective (Mld) have been shown to be involved in the control of ecdysone production during early larval stages in *Drosophila* [56]. Orthologues of *Drosophila kni* and *vvl* have been identified in *Tribolium*, showing conserved functions during embryogenesis [57,58]. Interestingly Tc-vvl has been shown to be a key factor coordinating ecdysteroid biosynthesis as well as molting in larval stages [59], supporting the idea that Tc-vvl also exerts similar function in the PG during early larval development. Nevertheless, further studies are required to confirmed the role of these genes and to elucidate ecdysone biosynthesis regulation in all larval transitions.

Why is then Egfr signaling specifically required during the larva-pupa transition? One possibility is that the peak of 20E required to trigger metamorphosis demands the growth of PG cells to increase the biosynthetic enzyme transcription. Interestingly, this process has recently been related to the nutritional state of the animal. In fact, it has been shown in *Drosophila* that surpassing a critical weight checkpoint that occurs at the onset of the last larval instar is required for the growth of the PG cells and the increase of ecdysone production [1,2,60,61], which implies that PG cells must reach a certain size to produce enough ecdysone to trigger metamorphosis. Similarly, in *Tribolium* exists a correlation between the attainment of a threshold size, defined as the mass that determines that the animal is in the last larval instar, and the upregulation of key metamorphic genes that induce the metamorphic transition [62]. Therefore, it is plausible that Egfr signaling is particularly required once the animal reaches the threshold size that determines the initiation of metamorphosis at the ensuing molt. Such activation will ensure the proper increase of ecdysone production to induce the formation of the pupa. In this sense, it is interesting to note that the highest expression of *Tc-Egfr* is detected at the last larval stage. Similarly, the expression of both ligands *Tc-spi* and *Tc-vn* are up-regulated during the late part of the last larval instar, supporting the fact that both ligands are required for Egfr signaling at this stage.

### *Tribolium* possesses two Egfr ligands

To date, a single Tc-Egfr ligand with high sequence similarity to *Drosophila* Dm-Spi had been identified in the genome of *Tribolium* probably due to inaccurate annotation of the beetle genome and the fact that its depletion in the embryo phenocopied *Tc-Egfr* knockdown. However, our *in silico* analysis has revealed the presence of a second Tc-Egf ligand, the neurogulin Tc-Vn. We have confirmed the identity of Tc-Vn in *Tribolium* by analyzing its protein sequence and its role during development. Phylogenetic analysis clustered Tc-Vn with its coleopteran orthologous proteins. Interestingly, based on the length of the phylogenetic arm, Tc-Vn presents a high range of change, probably due to its minor role on oogenesis and embryogenesis that has reduced the evolutive pressure on Tc-Vn ligand. In contrast, a lower divergence of Tc-Spi is observed, clustering closely to the *Drosophila* Dm-Egf ligands Dm-Krn and Dm-Spi. This observation is consistent with Tc-Spi exerting a key role during early development as main activator of Tc-Egfr signaling.

Our results indicate that both Egf ligands Tc-Vn and Tc-Spi function redundantly in the control of larval-pupal transition in *Tribolium* similarly to what occurs in *Drosophila*. In contrast, the requirement of each ligand is specific for the development of different structures during the pupal stage. Thus, Tc-Vn is required for the formation of wings and elytra, whereas Tc-Spi is responsible of the proper development of gin traps and the urogomphi as well as the mouthpart. In addition, depletion of either *Tc-spi* or *Tc-vn* produce distinct defects on the distal part of the leg and the antenna suggesting again a different role on the activation of Egfr signaling in these structures. This different ligand requirement is similar to what has been described in *Drosophila*. Thus, Dm-Vn is required for wing identity during early imaginal disc development [63], whereas later on, the combination of Dm-Spi and Dm-Vn are necessary for vein formation [64]. Similarly, both ligands Dm-Spi and Dm-Vn control leg patterning and growth [37], although Dm-Vn seems to have a predominant role in the formation of the tarsus, the most distal part of the leg [40,65]. In addition, our data also show that the activation of the Tc-Egfr pathway in oogenesis relay entirely on Tc-Spi. In this context, the similarity of Tc-Spi to the Dm-Egf *Drosophila* ligand Dm-grk [35], a dedicated ligand specific of the ovaries, might explain this exclusive requirement in this process in *Tribolium*. Therefore, our results confirmed that Egfr signaling is activated by at least two different ligands in most of the holometabolous insects in morphogenetic processes. The different expression and activity of those ligands probably has favored the co-option of Egfr signaling in different processes, from oogenesis to the biosynthesis of ecdysone, contributing to the generation of new organ and shapes along evolution.

## MATERIALS AND METHODS

### Tribolium castaneum

The enhancer-trap line pu11 of *Tribolium* (obtained from Y. Tomoyasu, Miami University, Oxford, OH) was reared on organic wheat flour containing 5% nutritional yeast and maintained at 29 °C in constant darkness.

### Conserved domains analysis and alignment of Vein sequences

Tc-Vn (TC014604) amino acid sequence was analyzed by different on line applications to detect its protein domains. Thus, peptide signal was predicted by SignalP 4.1 software that detect the presence and the location of signal peptide cleavage site [66]. Conserved sequences of EGF-like and IG-like domains were detected by NCBI’s Conserved Domain Database [67]. PEST region was predicted using EPESTFIND online software that detects PEST motifs as potential proteolytic cleavage sites. Multiple alignment was performed by ClustalW software [68]. Vein sequence of *D. melanosgaster* (CG10491), *Aedes aegypti* (XP_021697922), *Operophtera brumata* (KOB67206), *Bombix mori* (XP_004925912), *Trichogramma pretiosum* (XP_023315843) and *Periophthalmus gracilis* (XP_020280346), which were used for the alignments, were obtained from GenBank database.

### Phylogenetic analysis of Tc-Vn

To understand the phylogenetic relationship of Vein proteins, amino acid sequences from Vein proteins were collected from different insect taxa, including that of *Tribolium* as well as from four primates species as an outgroup (Supplementary Table 1) and aligned using MAFFT [69] v7.299b L-INS-I with 1000 iterations. Ambiguously aligned positions were trimmed using trimAlv14 [70], with the automated 1 algorithm. The best substitution model for phylogenetic inference was selected using IQ-TREE [71] with the TESTNEW model selection procedure and following the BIC criterion. The LG substitution matrix with a 4-categories discrete Γ distribution and allowing for invariant sites was selected as the best-fitting model. Maximum likelihood inferences were performed with IQ-TREE, and statistical supports were drawn from 1,000 ultrafast bootstrap values with a 0.99 minimum correlation as convergence criterion, and 1,000 replicates of the SHlike approximate likelihood ratio test.

### Quantitative real-time reverse transcriptase polymerase chain reaction (qRT-PCR)

Total RNA from individual larva of *Tribolium* was extracted using the GenElute™ Mammalian Total RNA kit (Sigma). cDNA synthesis was carried out as previously described [72,73]. Relative transcript levels were determined by quantitative real-time PCR (qPCR), using Power SYBR Green PCR Mastermix (Applied Biosystems). To standardize the quantitative real-time RT-PCR (qPCR) inputs, a master mix that contained Power SYBR Green PCR Mastermix and forward and reverse primers was prepared to a final concentration of 100 µM for each primer. The qPCR experiments were conducted with the same quantity of tissue equivalent input for all treatments, and each sample was run in duplicate using 2 µl of cDNA per reaction. As a reference, same cDNAs were subjected to qRT-PCR with a primer pair specific *Tribolium* Ribosomal Tc-Rpl32. All the samples were analyzed on the iCycler iQReal Time PCR Detection System (Bio-Rad). Primer sequences used for qPCR for *Tribolium* are:

Tc-EGFR-F: *5’*- TCACGAGCATGTGGTTATGAT -3’

Tc-EGFR-R: 5’- CTCATTCTCGAGCTGGAAGT -3’

Tc-Pnt-F: 5’- AGAGTTCTCCCTCGAATGCAT -3’

Tc-Pnt-R: 5’- TCTGCAACAACTCCAAGTGCT -3’

Tc-Spi-F: 5’- AACATCACATTCCACACGTAC -3’

Tc-Spi-R: 5’- TCTGCACACTCGCAATTGTAT -3’

Tc-Vn-F: 5’- GAAGTCCAAGACACACAACTC -3’

Tc-Vn-R: 5’- CTTGTATAGGTACCAGGTGTCT-3’

Tc-E93-F: 5’-CTCTCGAAAACTCGGTTCTAAACA-3

Tc-E93-R: 5’-TTTGGGTTTGGGTGCTGCCGAATT-3’

Tc-HR3-F: 5’- TCACAGAGTTCAGTTGTAAACT *-3’*

Tc-HR3-R: *5’-* TCTCGCTGCTTCTTCGACAT -3’

Tc-E75-F: 5’- CGGTCCTCAATGGAAGAAAA *-3’*

Tc-E75-R: *5’-* TGTGTGGTTTGTAGGCTTCG -3’

Tc-Br-C-F: 5’- TCGTTTCTCAAGACGGCTGAAGTG *-3’*

Tc-Br-C-R: *5’-* CTCCACTAACTTCTCGGTGAAGCT-3’

Tc-Phm-F: 5’-TGAACAAATCGCAATGGTGCCATA *-3’*

Tc-Phm-R: *5’-*TCATGGTACCTGGTGGTGGAACCTTAT -3’

Tc-Rpl32-F: 5’- CAGGCACCAGTCTGACCGTTATG *-3’*

Tc-Rpl32-R: *5’-* CATGTGCTTCGTTTTGGCATTGGA -3’

### Larva and adult RNAi injection

Tc-Egfr dsRNA (IB_00647), Tc-Pnt dsRNA (IB_02295), Tc-Spi dsRNA (IB_03555) and Tc-Vn dsRNA (IB_05654) were synthesized by the Eupheria Biotech Company. A concentration of 1 µg/µl dsRNA was injected into 50 larvae from penultimate instar and antepenultimate instar and 40 female adults in the abdominal body cavity laterally to avoid damaging genitals as previously described [74]. Both *Tc-Egfr*^*RNAi*^ and *Tc-Vn*^*RNAi*^ injected females were crossed with wild-type males in order to obtain *knockdown* embryos.

### Microscopy analysis

All ovaries were dissected in PBS, fixed with 3.7% formaldehyde/PBS and incubated with Phalloidin. The stained ovaries were mounted in Vectashiled with DAPI for epifluorescence microscopy. Dissected pupal appendages, were fixed with 3.7% formaldehyde/PBS and mounted in Vectashield with DAPI to visualized the nucleus. All pictures were obtained with AxioImager.Z1 (ApoTome 213 System, Zeiss) microscope, and images were subsequently processed using Fuji and Adobe photoshop.

## ACKNOWLEDGEMENTS

Support for this research was provided by the Spanish MINECO (grants CGL2014-55786-P and PGC2018-098427-B-I00 to D.M. and X.F-M.) and by the Catalan Government (2014 SGR 619 to D.M. and X.F-M.). The research has also benefited from FEDER funds. S.C. is a recipient of a pre-doctoral research Grant from the MINECO.

## AUTHOR CONTRIBUTIONS

Conception and design of the project was done by S.C., D.M. and X.F-M. S.C. and X.F-M performed the experiments. The analysis of the data was conducted by S.C, D.M. and X.F-M. X. F-M. wrote the manuscript and S.C. and D.M. revised the manuscript. All authors approved the final manuscript.

